# Visual mismatch negativity to disappearing parts of objects and textures

**DOI:** 10.1101/486191

**Authors:** István Czigler, István Sulykos, Domonkos File, Petia Kojouharova, Zsófia Anna Gaál

## Abstract

Visual mismatch negativity (vMMN), an event-related signature of automatic detection of events violating sequential regularities is traditionally investigated to the onset of frequent (standard) and rare (deviant) events. In a previous study [4] we obtained vMMN to vanishing parts of continuously presented objects (diamonds with diagonals), and we concluded that the offset-related vMMN is a model of sensitivity to irregular partial occlusion of objects. In the present study we replicated the previous results, but to test the object-related interpretation we applied a new condition with a set of separate visual stimuli: a texture of bars with two orientations. In the texture condition (offset of bars with irregular vs. regular orientation) we obtained vMMN, showing that the continuous presence of objects is unnecessary for offset-related vMMN. However, unlike in the object-related condition, reappearance of the previously vanishing lines also elicited vMMN. In a formal way reappearance of the stimuli is an event with probability 1.0, and according to the results, object condition reappearance is an expected event. However, offset and onset of texture elements seems to be treated separately by the system underlying vMMN. As an advantage of the present method, the whole stimulus set during the inter-stimulus interval saturates the visual structures sensitive to stimulus input. Accordingly, the offset-related vMMN is less sensitive to low-level adaptation difference between the deviant and standard stimuli.

## 1. Introduction

The visual information processing system is sensitive to events violating the regularity of stimulus sequences, even if the events are unrelated to the ongoing task (unattended). The automatic detection of violating regularities can be revealed by the visual mismatch negativity (vMMN) components of event-related brain potentials (ERPs). VMMN is the difference between the ERPs elicited by the deviant events and the ERPs to the regular ones. VMMN is elicited by deviant visual features (color, orientation, movement direction, etc.), object-related deviancies, facial emotions, handedness, numerosity, sequential regularities, familiarity, language-related and other deviances, etc. (for reviews see [1, 2, 3]).

In our previous study [4] we obtained vMMN to the offset of irregularly vanishing part of objects. In particular, diamonds with diameters were presented during the interevent interval. From time to time two parallel sides of the diamonds disappeared. One of the parallel sides disappeared infrequently, the other pair disappeared frequently. Importantly, diamonds were unrelated to the ongoing tracking task. VMMN, as a difference potential between those elicited by the infrequent and frequent offset emerged over the occipital location within the 120-202 ms range. However, no vMMN appeared after the reappearance of the whole object. We interpreted our result as showing that the infrequent occlusion of the represented objects elicited vMMN, whereas the reappearance of the object was a predicted event, and accordingly these events did not elicit vMMN. This interpretation is in accord with a prevailing theory of auditory MMN and vMMN. The predictive coding theory considers the mismatch potentials as errors signals. The memory representation of the frequent (standard) stimuli generates an expectation about the likely properties of future events. In case of match between the input and the expected representation (i.e., without new information) the perceptual system may ignore event. Further processing only occurs when there is discrepancy between the input and the expectancy. The mismatch components are signatures of the mutual adjustment between the input and the expected events only. According to the predictive coding view, reappearance of the whole pattern (i.e. an event with 1.0 probability) does not elicit vMMN [2, 3, 5, 6]. Furthermore, this interpretation of the previous study [4] was closely connected to object-related representation, because we considered that the environmental model consisted of the representation of the whole diamonds. The aim of the present study was to replicate this result, and investigate the object-related aspect of our interpretation. On this end beside the object-related condition, in the inter-event period we presented unconnected bars with two orientations (texture condition). One set of bars with a particular orientation vanished infrequently, the other frequently. We hypothesized, that without the object-related representation stimulus offset does not elicit vMMN, but stimulus onset, as an orientation-related deviancy elicits vMMN.

It is important to note that the offset stimulation has a particular advantage. While the stimuli are present during the inter-event interval, these stimuli saturate the low-level input structures. Therefore the ERPs to deviant vs. standard difference are less susceptible to stimulus-specific adaptation, therefore offset-related vMMN can be considered as deviant-related additional activity (genuine vMMN; [7, 8]).

## 2. Materials and methods

### 2.1 Participants

Twenty adults participated in the study. All of them had normal or corrected-to-normal vision (at least 5/5 in a version of the Snellen charts). No one reported any neurological or psychiatric diseases. They were paid for their participation. One of the participants had an unusually noisy ERP, and another participant’s ERP was dominated by alpha activity. Therefore, the results were calculated for the remaining 18 participants (10 females, mean age: 22.1 years, SD: 2.3 years). Participants were paid for their contributions. Written informed consent was obtained from the participants before the experimental procedure. The study was approved by the United Ethical Review Committee for Research in Psychology (Hungary).

### 2.2 Stimuli and procedure

The experimental stimuli of the object condition and other aspects of the study were identical to our previous study [4]. As a summary, events were presented on a 19-in CRT monitor (Flatron 915 FT Plus, 75 Hz refresh rate) from a 1.4 m distance using the Cogent 2000 MATLAB toolbox. Figure 1 demonstrates the task-related and vMMN-related stimuli in the two conditions and the stimulus sequence.

**Figure 1.**
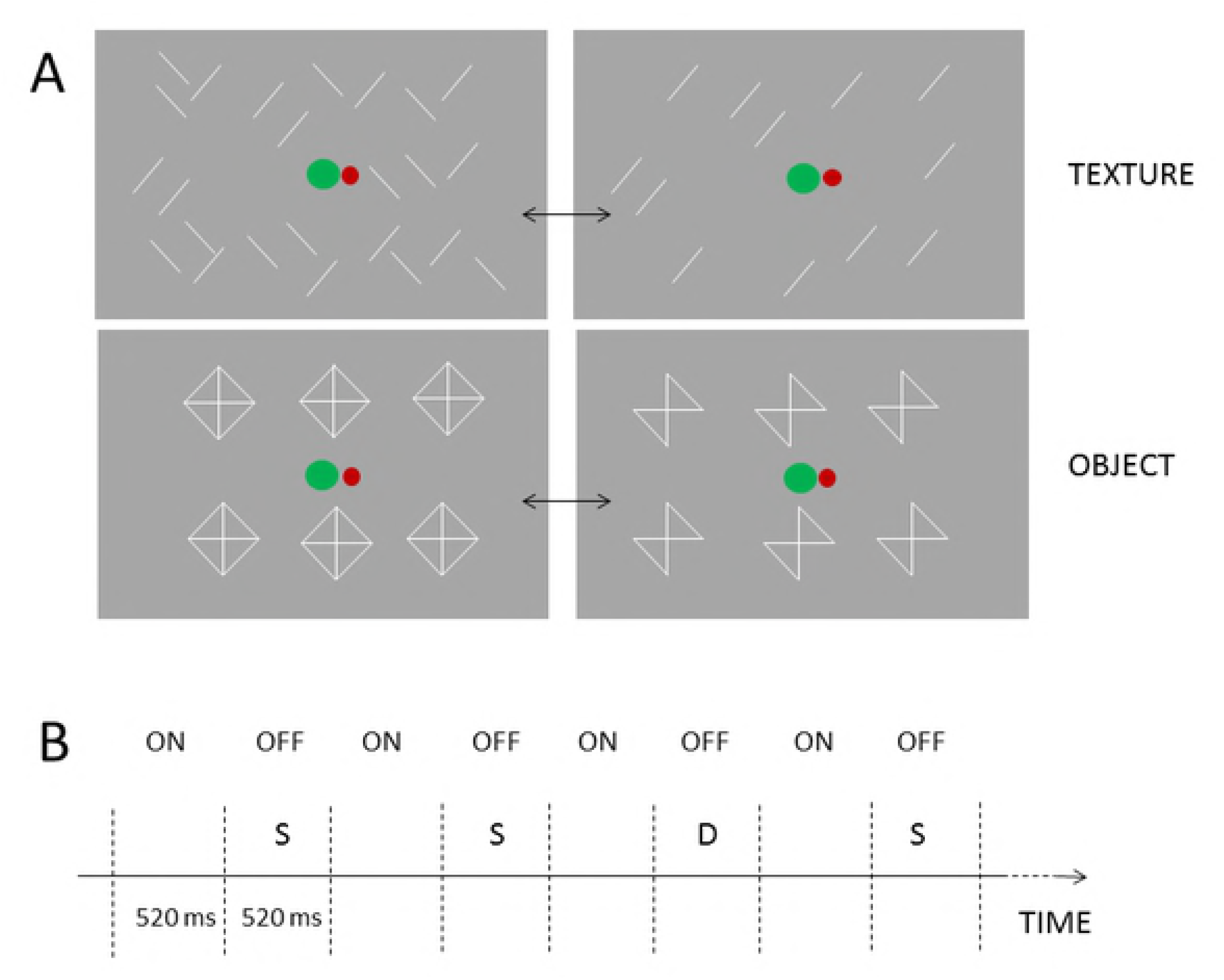
Stimuli and stimulus sequences in the texture and object conditions. A: An example of the stimulus field. The vanishing stimuli were either the 45o or the 135o bars. Both orientations were standard and deviant. B: the outline of the stimulus sequences (in both the onset and offset stimuli a +/− 40 ms range was presented around the 520 ms mean value). The green and red dots are the stimuli of the tracking task.

The task-relevant stimuli appeared on the central area of the screen and consisted of two disks. The red disk served as a fixation point, the green disk made horizontal random motion around the red disk. The task was to keep the green disk as close to the center of the red disk as possible with the left and right arrows of a keyboard. Error occurred when the distance of the two disks exceeded 1.1 degrees. Performance (the sum of errors in one block) was reported on the screen at the end of each block. Behavioral data were defined as the number of occasions when the ball left the target area. The performance of the two conditions was compared in a t-test.

The vMMN-related irrelevant stimuli appeared around the task-relevant stimuli. In the *Object condition*, diamonds and diamonds without two of their parallel lines appeared alternately. Six identical objects (75.5 cd/m^2^) were presented (in a 2 row by 3 against a medium-gray background (20.1 cd/m^2^). There was no inter-stimulus interval between these patterns. In the offset events either the two 45-degree sides or the two 135-degree sides of the diamonds were omitted. These two patterns were presented in oddball sequences, with either the left-tilted or the right-tilted version as deviant (p=0.2). In one block there were 95 offset events, 76 standard bow ties, and 19 deviant ones. According to the reverse control principle, both the left- and right-tilted bow ties served as deviant and standard (6 sequences for each). Altogether, 570 stimuli were presented in each deviant-standard direction. The stimulus duration of all three patterns was 520 ms (with +/− 40 ms jitter in 13.3 ms steps).

In the *Text condition,* there were oblique lines with 45-degree and 135-degree orientations. The lines were randomly dispersed within the stimulus field, but the number of tilted lines, the size of the lines and the luminances were equal to those in the *Object* condition, and in all other respects, the two conditions were identical. Figure 1A demonstrates the screen of task-related and vMMN-related stimuli in the two conditions, and Figure 1B shows the stimulus sequence.

### 2.3 EEG recording, ERP acquisition and measurement

EEG was recorded with a Neuroscan recording system (SynampsRT amplifier, Compumedics Abbotsford Ltd, Australia, EasyCap, Advanced Medical Equipments Ltd, Horsham, UK; Ag/AgCl electrodes, DC-200 Hz, sampling rate: 1000 Hz). Thirty-eight electrode locations were used, in accordance with the extended 10–20 system. The ground electrode was placed on the forehead. An electrode on the tip of the nose served as a reference. HEOG and VEOG were recorded with bipolar configurations between two electrodes placed laterally to the outer canthi of the two eyes or above and below the left eye, respectively.

The EEG signal was analyzed with a MATLAB script developed in our lab. First, it was filtered offline with a noncausal Kaiser-windowed finite impulse response filter (low pass: 30; high-pass: 0.1 Hz). Epochs of 600 ms (including 100 ms prestimulus interval serving as baseline) were extracted for all deviants and for those standards that immediately preceded the deviants. Epochs with larger than 100 μV or smaller than 2 μV voltage change were considered artifacts and rejected from the further processing. ERPs were calculated by averaging the extracted epochs. According to the reverse control principle, epochs from both experimental (oddball and reverse) sequences were entered into the averaging process.

Event-related potentials were averaged separately for the two conditions (object and text), and within the conditions for the two events (offset and onset) and for the two probabilities (deviant, standard). Only those ERPs to the standard stimuli were included in the averaging that appeared before a deviant. The number of averaged epochs was 3828 and 3827 for deviants and last standards which is 84% of all epochs.

On the basis of the results of our previous study [4] we calculated an occipital ROI (O1, Oz, O2) from the deviant minus standard difference potentials. In the previous study vMMN emerged at the occipital locations within the 220-202 ms range, therefore in the present study we calculated the mean activity within this range. VMMN amplitudes were compared in a two-way ANOVA with factors of *Condition* (object, texture) and *Event* (offset, onset).

To control the reliability of difference between the ERPs to the deviant and standard, within the possible vMMN range we calculated series of t-tests over the 100-300 ms range at O1, Oz and O2 electrodes on the deviant minus standard difference potentials (difference from zero). As a criterion of 25 consecutive t-values (25 ms) were significant (p<0.05) at least over two locations. We obtained significant values within 116-178 ms, 139-195 ms and 155-208 ms ranges (i.e., 62 ms, 56 ms and 53 ms) for object offset, texture offset and texture onset, respectively.

As unexpected findings, in comparison to the standard stimuli, following vMMN, both offset and onset deviants elicited posterior positivity. Furthermore, over the anterior locations positive difference potentials emerged, and these positivities were larger for the offset stimuli. We measured the peak latency and the amplitude values of these positive differences in the posterior and anterior ROIs (O1, Oz, O2 and F3, Fz and F4, respectively). Latencies were measured as the largest positive component within 200-300 ms¸ and amplitudes were measured as the mean activity of this range. These measures were analysed in ANOVAs with factors of *Condition* (object, text) and *Event* (offset, onset).

To compare the ERPs to stimulus onset and offset on the exogenous activity, we measured the latencies and amplitudes of the posterior exogenous negative component (N1) on the occipital ROI (O1, Oz, O2). N1 component was identified in the 120-200 ms window as the highest negative-going deflection, and its latency was measured on the standard stimuli. Amplitudes were measured as the means of a +/− 5 ms range around the group average. The amplitudes and latencies were compared in ANOVAs with factors of *Condition* (object, texture) and *Events* (offset, onset). In the ANOVAs effect size was calculated as partial eta squared (η_p_^2^).

## 3. Results

### 3.1 Behavioral results

Performance (errors) was characterized by the number of cases when the distance of the two discs exceeded 1.1 deg. Performance was fairly high, and the group average of errors were 5.56 (SD=1.97) and 7.17 (SD=4.23) in the object and texture conditions, respectively. In a t-test, the difference was not significant.

### 3.2 Event-related potentials

As Figure 2 and 3 shows, deviant object offset, texture offset and texture onset elicited a negative deviant minus standard posterior difference potential, but object onset did not elicit posterior negativity. To replicate the results [4] we calculated vMMN amplitude within the range of significant difference of the previous study (120-202 ms). In an ANOVA with factors of *Condition* and *Event*. We obtained significant main effect of *Event*, F(1,17)=5.23, p=0.035, η_p_^2^=0.24, and interaction F[1,17]=73.28, p=0.029, η_p_^2^=0.25). Following the negative difference potentials, for the deviant offset events positivities emerged over the posterior and anterior locations within the 200-300 ms range (Table 1). We conducted separate ANOVAs for the posterior (O1, Oz, O2) and anterior (F3, Fz, F4) ROIs with factors of *Condition* and *Event*. According to the ANOVA the main effect of *Event* was significant, F(1,17)=8.39, p=0.010, η_p_^2^=0.33. In a similar ANOVA for the anterior positivity the main effect of *Event* was also significant, F(1,17)=8.26, p=0.011, η_p_^2^=0.30. Table 1 shows the amplitude values of the negativity.

**Figure 2.**
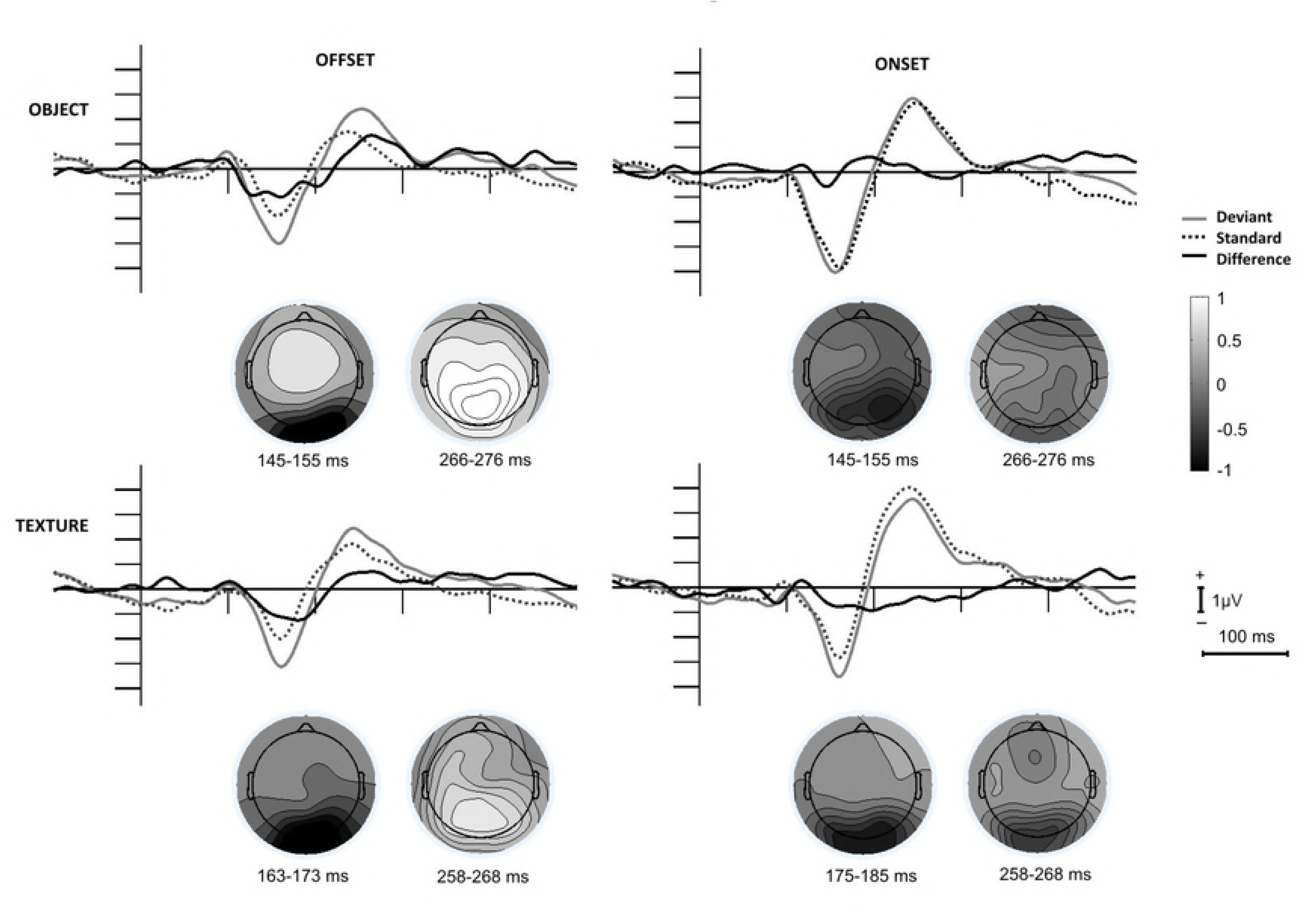
Event-related potentials and difference potentials at the posterior (occipital) ROI to stimulus offset and onset events in the Object and Texture conditions. The scalp distributions are calculated for the ranges with significant deviant minus standard differences.

**Figure 3.**
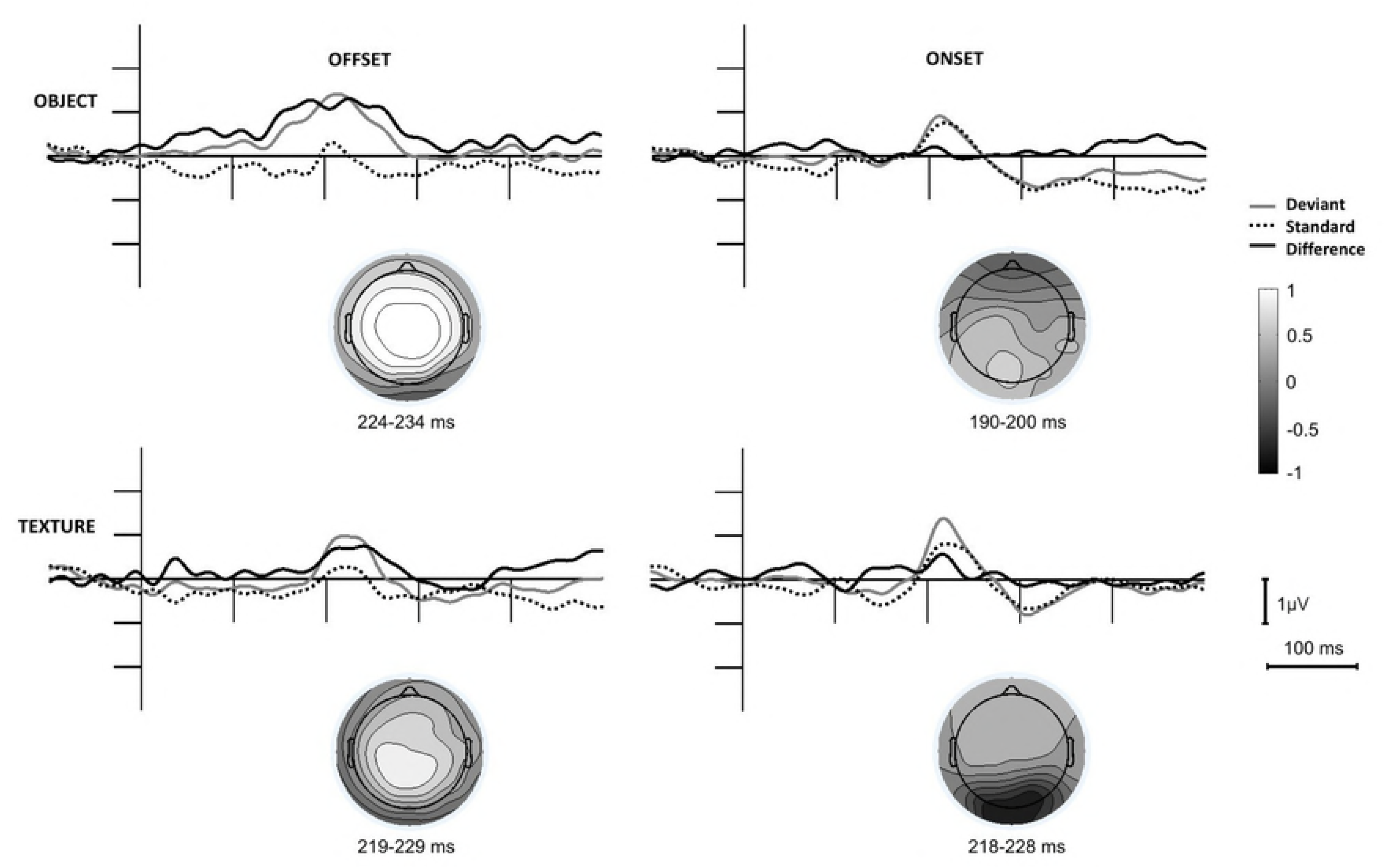
Event-related potentials and difference potentials at the anterior (frontal) ROI to stimulus offset and onset events in the Object and Texture conditions. The scalp distributions are calculated for the ranges with significant deviant minus standard differences.

As Figures 2 and 3 show, positive difference potentials emerged over the posterior and anterior locations. To explore the appearance of the positivities in the two conditions to the two events, we conducted ANOVAs on the mean amplitudes within the 200-300 ms latency range. For both ROIs the *Event* main effect was significant: F(1,17)=8.38, p=0.010, η_p_^2^=0.33 for the occipital (O1, Oz, O2) ROI, F(1,17)=9.75, p=0.010, η_p_^2^=0.33 and F(1,17)=8.26, p=0.011, η_p_^2^=0.30 for the anterior ROI (F3, Fz, F4), respectively, indicating larger positivity to the offset events.

To compare the ERPs in the texture and object conditions to the onset and offset events, ANOVAs with factors of *Condition* and *Event* were calculated for the peak latency and the mean amplitude values (+/− 5 ms around the group average). Latency values were fairly similar, 160 ms, 157 ms, 158 ms and 162 ms for object offset, object onset, texture offset and texture onset, respectively. Accordingly, neither the main effects, nor the interaction were significant. As Table 1 shows, onset events elicited larger N1 than offset events. In the ANOVA the *Condition* main effect was significant, F(1,17)=22.31, p<0.001, η_p_^2^=0.57. According to the significant interaction, F(1,17)=20.71, p<0.001, η_p_^2^=0.55, the difference was due to the larger N1 to the object onset.

**Table 1.**
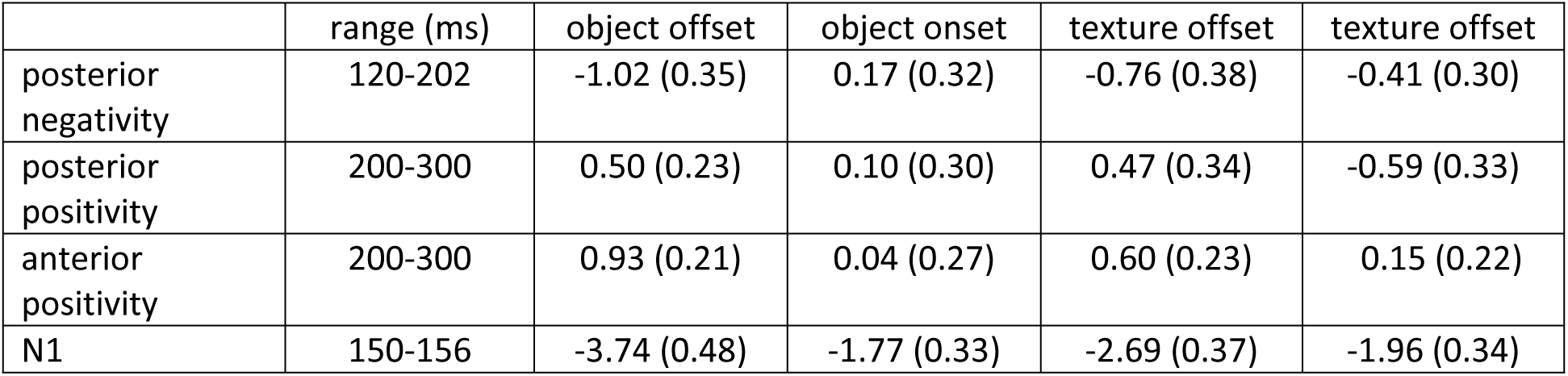
Amplitude values (µV) of the posterior negative difference potential (vMMN), the positive and anterior positivities and the N1 components (standard error of mean in parenthesis).

|  | range (ms) | object offset | object onset | texture offset | texture offset |
| --- | --- | --- | --- | --- | --- |
| posterior negativity | 120-202 | -1.02 (0.35) | 0.17 (0.32) | -0.76 (0.38) | -0.41 (0.30) |
| posterior positivity | 200-300 | 0.50 (0.23) | 0.10 (0.30) | 0.47 (0.34) | -0.59 (0.33) |
| anterior positivity | 200-300 | 0.93 (0.21) | 0.04 (0.27) | 0.60 (0.23) | 0.15 (0.22) |
| N1 | 150-156 | -3.74 (0.48) | -1.77 (0.33) | -2.69 (0.37) | -1.96 (0.34) |

## 4. Discussion

On the basis of the object-related representation of environmental events coding we expected vMMN to object offset, but we were uncertain whether the offset of visual textures elicit vMMN. Furthermore, we expected no onset-related vMMN with object onset deviancy. According to the results both object and texture offset elicited vMMN. Concerning the reappearance of objects there were no detectable ERP difference between the onset after the frequently and infrequently vanishing lines of the diamonds. In other words, in the object condition we did not register vMMN. This result replicated our previous finding [4]. However, emergence of vMMN to texture offset and onset requires the revision of the view suggested by Sulykos and colleagues [4]. We claimed that the memory system underlying vMMN represented objects as wholes (Gestalts), and in the study the offset stimuli was a model of partial occlusion. Therefore VMMN emerged when frequent occlusions were replaced by rare ones. Furthermore, reappearance of the object, irrespective of the previous (deviant or standard) offset was a fully predictable event, therefore this event did not elicit vMMN. However, as the offset-related vMMN of the texture condition of the present study shows, vanishing of particular bar orientations were sufficient for eliciting vMMN. Importantly, there was an obvious difference between the object and texture conditions, i.e., appearance of onset-related vMMN in texture condition. To preserve an aspect of the object-related representation, we claim that in the object condition the system underlying vMMN treated the offset and onset events as units (disappearance and reappearance of parts of the objects). However, the system underlying vMMN treats texture offset and onset separately, i.e., rare vs. frequent offset of particular line orientations, and rare vs. frequent onset of particular line orientations. In other words, whereas the representation of the object survived the offset period, texture onset and offset were treated as separate events. This explanation preserves the notion that vMMN is a surprise-related component elicited by non-reinforced predictions [e.g. 4, 9, 10, 11, 12, 13, 14], even if in a formal sense, onset is a fully expected event in both conditions.

Posterior positivity following the vMMN appeared in previous studies [16, 17]. In the present study this positivity appeared only to the offset events. Similarly, anterior positivity appeared in some studies [7, 18] to deviant stimuli. However, connection of these positivities to the processes underlying vMMN and their functional significance is unclear. Furthermore, some recent studies reported positive mismatch responses emerged in later latency ranges [9, 19]. Due to the lack of a priory expectation, as a speculative explanation, the positivities are connected to a further processing of the more salient offset stimulation; in this case anterior structures are involved in the processing of deviant events. The predictive coding view [5] is capable of explain these ERP effects as a modification of the environmental model. However, relations between vMMN and the subsequent positivities require further research.

Onset events usually elicit ERPs with larger amplitudes than offset events [20, 21]. We obtained similar results. Onset-related N1 was larger in the object condition. While the reappearing bars were similar in the two conditions, we have no post-hoc explanation for this unexpected result.

In conclusion, offset stimuli after a longer onset period potentially saturated the input-related visual structures. However, infrequently vanishing stimulus elements elicited the signature of automatic deviance detection, the visual mismatch negativity. In comparison, to textures consisted of unconnected bars the memory system underlying vMMN predicted the reappearance of Gestalt-like stimuli (objects), and stimulus onset of the objects did not elicit vMMN. As a tentative suggestion, in a visual scene disappearance can be a more salient event than reappearance, and the more salient event may lead to further processing, as indicated by both posterior and anterior activity.

## Acknowledgments

This research was supported by National Research, Development and Innovation Office (NKFIH), Grant 119587.

